# Targeted delivery of rhodopsin’s assembled core is required for outer segment extension in mouse rod photoreceptors

**DOI:** 10.1101/2024.12.23.630147

**Authors:** Jorge Y. Martínez-Márquez, Sandy Hua, Andreea M. Beu, Christopher B. Stein, Jillian N. Pearring

## Abstract

Vertebrate vision in dim-light environments is initiated by rod photoreceptor cells that express the photopigment rhodopsin, a G-protein coupled receptor (GPCR). To ensure efficient light capture, rhodopsin is densely packed into hundreds of membrane discs that are tightly stacked within the rod-shaped outer segment compartment. Along with its role in eliciting the visual response, rhodopsin serves as both a building block necessary for proper outer segment formation as well as a trafficking guide for a few outer segment resident membrane proteins. An interesting aspect of rod homeostasis is that mutations that affect the localization of rhodopsin to the outer segment result in photoreceptor degeneration. In this study, we focus on determining the necessary properties of rhodopsin’s cytosolic C-terminus required for either proper outer segment trafficking and/or the capacity to extend the rudimentary outer segment in rhodopsin knockout (RhoKO) rods. We find that the well-described C-terminal QVAPA outer segment targeting motif also plays a role in endoplasmic reticulum (ER) exit and is necessary for outer segment elongation. Even minor rhodopsin mislocalization prevents efficient outer segment elongation suggesting that QVAPA-targeted delivery of rhodopsin drives outer segment extension. We identify that rhodopsin’s core, helix-8, and accessible QVAPA targeting motif are the minimal requirements to extend the rudimentary outer segment in RhoKO rods. Our findings provide useful insights into rhodopsin’s molecular features needed for outer segment delivery and subsequent elongation of this membrane-rich compartment.

## INTRODUCTION

The retina lining the back of the eye contains photoreceptor cells that detect light. These photoreceptors have a modified primary cilium, called the outer segment, a cylindrical organelle that contains hundreds of stacked disc-shaped membranes. In rod photoreceptors, disc membranes are filled with rhodopsin, a GPCR that initiates the visual response to light (1). Rhodopsin is initially synthesized and trafficked through the ER and Golgi complex before being delivered via intracellular vesicle traffic to the apical membrane of the inner segment (2, 3). From here, rhodopsin travels through the connecting cilium to the outer segment where it is densely packed into nascent disc membranes. Currently, more than 150 variants in rhodopsin have been identified in human patients with retinitis pigmentosa, accounting for 30-40% of all cases (4, 5). Mutations that cause a defect in the trafficking and delivery of rhodopsin to the outer segment result in some of the most severe cases of autosomal dominant retinitis pigmentosa (6, 7). Understanding how rhodopsin is trafficked to the outer segment is imperative to develop future therapeutic targets for these patients.

As a prototypical GPCR, rhodopsin is a 7 transmembrane (TM) domain protein with an extracellular N-terminus and a cytosolic C-terminus. Rhodopsin’s extracellular N-terminus is glycosylated at two asparagine residues (N2 and N15) (8, 9) and mutations that affect the glycosylation result in photoreceptor degeneration in humans and mice (10, 11). The exact role of glycosylation remains unknown but is postulated to be involved in intrinsic folding, secretory processing, and forming intradiscal interactions that “Velcro” discs flat (10, 12). Rhodopsin’s core structure composed of 7-TM helices forms the binding pocket for the chromophore, 11-*cis*-retinal, which isomerizes to all-trans-retinal upon energy transfer from photons of light inducing rhodopsin conformational change and activation (13, 14). The core structure of rhodopsin is then followed by an eighth amphipathic α-helix (Helix-8, H8) and two palmitoylated cysteines (CC anchor) that help position H8 parallel to the membrane. Rhodopsin’s H8 has been implicated in binding to 11-*cis*-retinal, G-protein activation and protein folding (15).

Structural and biochemical studies have shown that rhodopsin molecules form homodimers (16, 17), and cryo-EM structural data shows that the interphase between TM1 and H8 mediates this dimerization (18). A recent study showed key residues in rhodopsin’s H8 form hydrophobic interactions with TM1 to stabilize the core structure of rhodopsin (19). While this study found that mutants disrupting these hydrophobic interactions cause mislocalization of rhodopsin to the inner segment, they also found that interchanging these key residues for each other retained the necessary hydrophobic interactions for proper outer segment localization. Finally, following the CC anchor, rhodopsin ends with a largely unstructured string of amino acids, 25 in mouse rhodopsin. This intracellular tail contains multiple sites for post-translational modification used to recruit proteins in the visual signaling pathway and encodes trafficking signals important for the delivery of rhodopsin to the outer segment (20–24).

Membrane proteins can ensure proper delivery to their site of action by encoding a targeting signal that interacts with specific transport partners as the proteins travel through the endomembrane system. The most studied outer segment targeting signal is rhodopsin’s C-terminal QVAPA targeting motif which is both necessary and sufficient for outer segment delivery (24, 25). Subsequent studies have found that while most transmembrane proteins encode targeting signals guiding their delivery to the outer segment (26–30), two outer segment resident proteins, GC-1 and PRCD, rely on interactions with rhodopsin for their delivery (31, 32).

Interestingly, untargeted membrane proteins exogenously expressed in rods strongly accumulate in the outer segment compartment with a minor fraction located in the plasma membrane. This localization pattern, previously coined the “default” trafficking pathway (33, 34), was postulated to occur because the outer segment is a membrane-rich organelle that does not undergo typical protein turnover. The outer segment undergoes continuous renewal by balancing the addition of new discs at the base with the phagocytosis of old discs from the distal tip by retinal pigmented epithelial cells (35, 36). Ongoing disc formation places a high demand on membrane transport toward the outer segment, which was thought to be driving the “default” pathway. However, it is generally believed that no resident protein relies on the “default” pathway for outer segment localization (29).

Rhodopsin is an integral building block of outer segment disc membranes. Loss of rhodopsin results in rods extending a small shapeless compartment filled with disorganized membranes from their connecting cilium (37–39). It was shown that these membranes largely preserve the full complement of outer segment resident proteins (31). The failure to elaborate a proper outer segment leads to the rapid degeneration of rod photoreceptors in RhoKO mice (37, 39). A previous study found that a Q334Ter rhodopsin truncation mutant was mislocalized to the inner segment and when in the presence of endogenous rhodopsin resulted in mislocalization of the latter (40). Moreover, Q334Ter rhodopsin expression could not properly elongate rod outer segments or rescue rod degeneration in RhoKO mice. These results suggest that specific targeting and efficient delivery of rhodopsin molecules to the outer segment is needed for outer segment formation and healthy rods. However, a more complete understanding of rhodopsin’s molecular features that are required for targeted delivery and subsequent outer segment elongation remain unknown.

In this study, we examined the key properties present in rhodopsin’s cytosolic C-terminus that ensure its proper trafficking, specific targeting and ability to build outer segments. We compared mutant rhodopsin constructs by analyzing their localization pattern in wildtype (WT) and RhoKO rods. In WT rods, rhodopsin mutant trafficking and targeting capacity was determined by the proportional mislocalization from the outer segment. In RhoKO rods, we examined localization patterns and the ability to extend the rudimentary outer segment for each rhodopsin construct we tested. Our data suggests that: 1) proper folding of the core rhodopsin molecule is needed for ER exit, 2) the QVAPA signal is required for outer segment targeting and formation, 3) minor mislocalization of rhodopsin prevents proper outer segment elongation and, 4) rhodopsin’s 7-TM core, helix-8 and unhindered access to QVAPA are required to extend an outer segment.

## RESULTS

### Pipeline to analyze membrane protein localization profiles of electroporated rod photoreceptors

We employed an *in vivo* electroporation technique to understand localization patterns of FLAG-tagged rhodopsin constructs in mouse rod photoreceptors. We began by establishing a standardized method to quantify the localization of a particular construct so that we could analyze data across animals and compare the localization profiles from different constructs. To do this we measured the fluorescence intensity present in the outer segment compartment, which was delineated by a soluble mCherry transfection marker co-electroporated with all our constructs. We used a soluble fluorescent marker because the photoreceptor outer segment is a membrane-rich compartment with very little cytosol, so the mCherry signal is primarily found in the inner segment, nuclear, and synaptic compartments. For a given image, the outer segment fluorescence signal of a particular construct was then divided by its total fluorescence in all rod compartments, which results in an OS/Total Intensity value for each image where 1.0 is fully localized to the outer segment and 0.0 is fully excluded from the outer segment.

We initially established the localization profile for a known outer segment protein by analyzing full-length FLAG-tagged rhodopsin (FLAG-Rho FL), which resulted in a mean OS/Total Intensity value of 0.908±0.061 (**Figure 1**). To establish the localization profile for an untargeted membrane protein we used an eGFP-mGluR1_TM1_ construct, previously shown to reach the outer segment through “default” trafficking in both WT mouse and frog rods (29, 33). This construct resulted in a mean OS/Total Intensity value of 0.485±0.173 (**Figure 1**). Finally, to determine the localization profile for a membrane protein that is excluded from the outer segment, we chose to express an eGFP-tagged transmembrane domain from a microsomal cytochrome b5 (eGFP-Cb5TM). This construct was previously shown to localize within the endoplasmic reticulum (ER) in frog rods (33) and therefore is not present within the outer segment compartment. The eGFP-Cb5TM construct resulted in a mean OS/Total Intensity value of 0.008±0.007 (**Figure 1**). These data establish the localization profile for a targeted, untargeted, and ER-retained membrane construct in WT rods.

**Figure 1:**
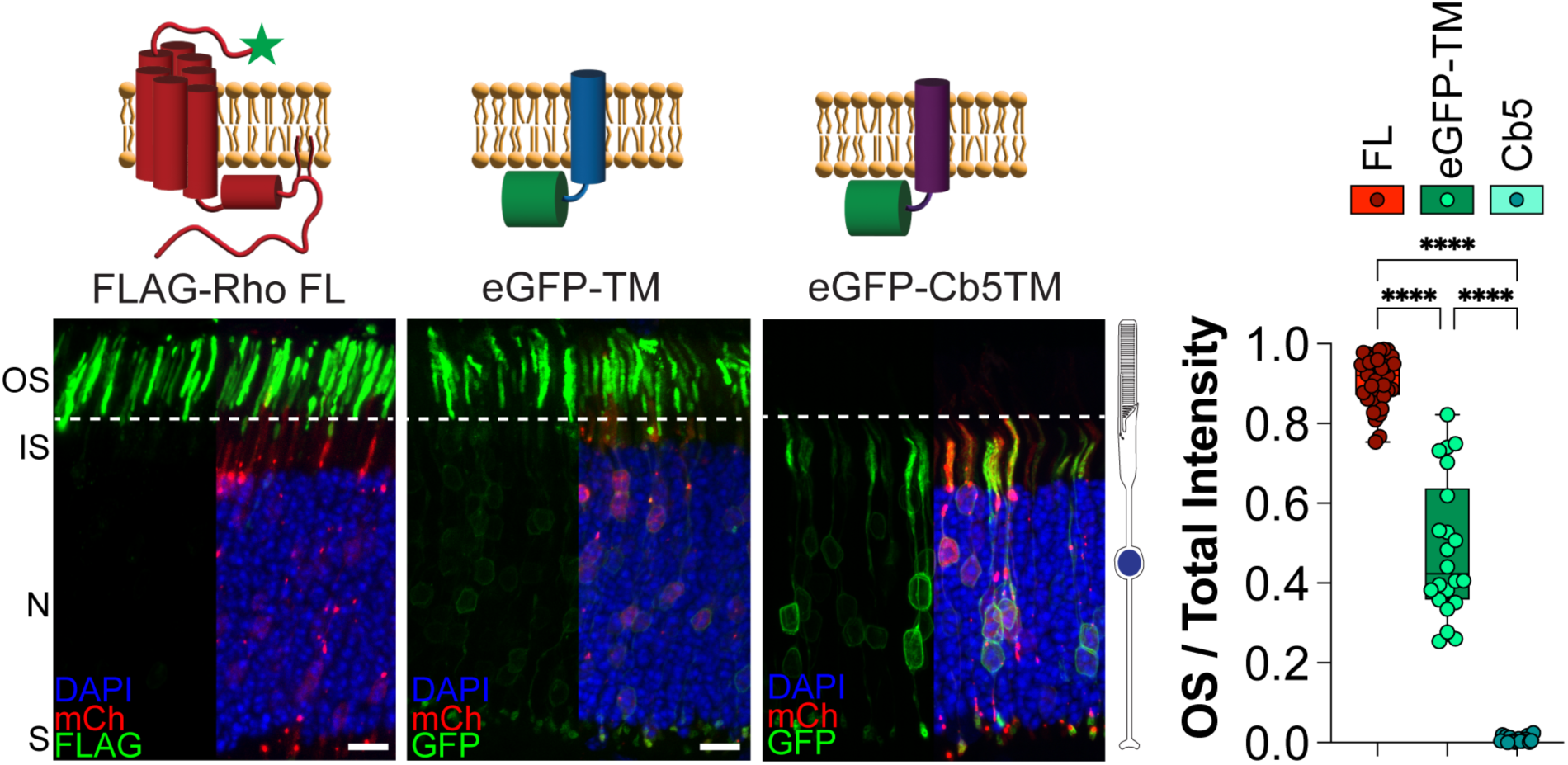
Localization profile of outer segment targeted, untargeted, and ER-retained membrane proteins in WT mouse rods. The following constructs were electroporated into WT mouse rods: FLAG-tagged full-length rhodopsin (Rho FL), eGFP-tagged transmembrane segment 1 from mGluR1 (eGFP-TM), eGFP-tagged transmembrane domain from microsomal cytochrome b5 (eGFP-Cb5TM). FLAG staining or GFP (green), mCherry (red) labels transfected rod cells and DAPI (blue) used to counter-stain nuclei. Scale Bar, 10 µm. To the right is a cartoon of a rod photoreceptor. Bar graph shows the quotient between the outer segment signal over the total signal for each construct. Each point represents a single image. FL n=12, 38 images; eGFP-TM n=7, 22 images; Cb5 n=6, 19 images. Here and in all figure panels: OS, outer segment; IS, inner segment; N, outer nuclear layer; S, synapses. Only half of DAPI and mCherry fluorescence is shown in the images to provide an unobstructed view of the green channel’s fluorescence. A cartoon diagram for each construct is shown on top of the representative image, with the FLAG tag depicted with a green star. All p-values can be found in **Sup Figure 1**.

### Determining the localization profiles of C-terminally truncated rhodopsin in WT mouse rods

We studied several rhodopsin truncation mutants to understand how regions within the cytoplasmic C-terminus of rhodopsin are involved in its proper delivery to the outer segment. We first expressed FLAG-tagged RhoΔ5 which lacks the outer segment QVAPA targeting signal and was previously shown to result in mislocalization of rhodopsin from the outer segment to the rest of the cell body (40). The RhoΔ5 construct was localized throughout the entire rod cell, from the basal synapse to the apical outer segment with a mean OS/Total Intensity value of 0.345±0.164 (**Figure 2A**). This profile reveals that there is significantly more of the RhoΔ5 construct present outside the outer segment than a typical untargeted membrane protein (see **Figure 1**), which is consistent with previously published data suggesting that there is a mistargeting signal residing downstream of the CC anchor in rhodopsin (324-329 amino acids) (20). We then expressed a FLAG-tagged RhoΔ25, which further truncates the C-terminus to the CC anchor, and found this construct was predominantly localized in the outer segment with a small fraction mislocalized to the cell body, showing a mean OS/Total Intensity value of 0.530±0.165 (**Figure 2A**). This localization profile is consistent with being an untargeted membrane protein as it is predominantly found within the membrane-rich outer segment. Finally, we expressed a FLAG-tagged RhoΔ38 that truncates the entire cytoplasmic C-terminus, including the amphipathic helix-8. We observed that RhoΔ38 could not reach the outer segment and appears to be held within internal membranes, likely the ER, as its mean OS/Total Intensity value was 0.016±0.017 (**Figure 2A**). Interestingly, each rhodopsin truncation revealed a unique localization profile with RhoΔ5 appearing mistargeted, RhoΔ25 appearing untargeted, and RhoΔ38 appearing ER-retained.

**Figure 2:**
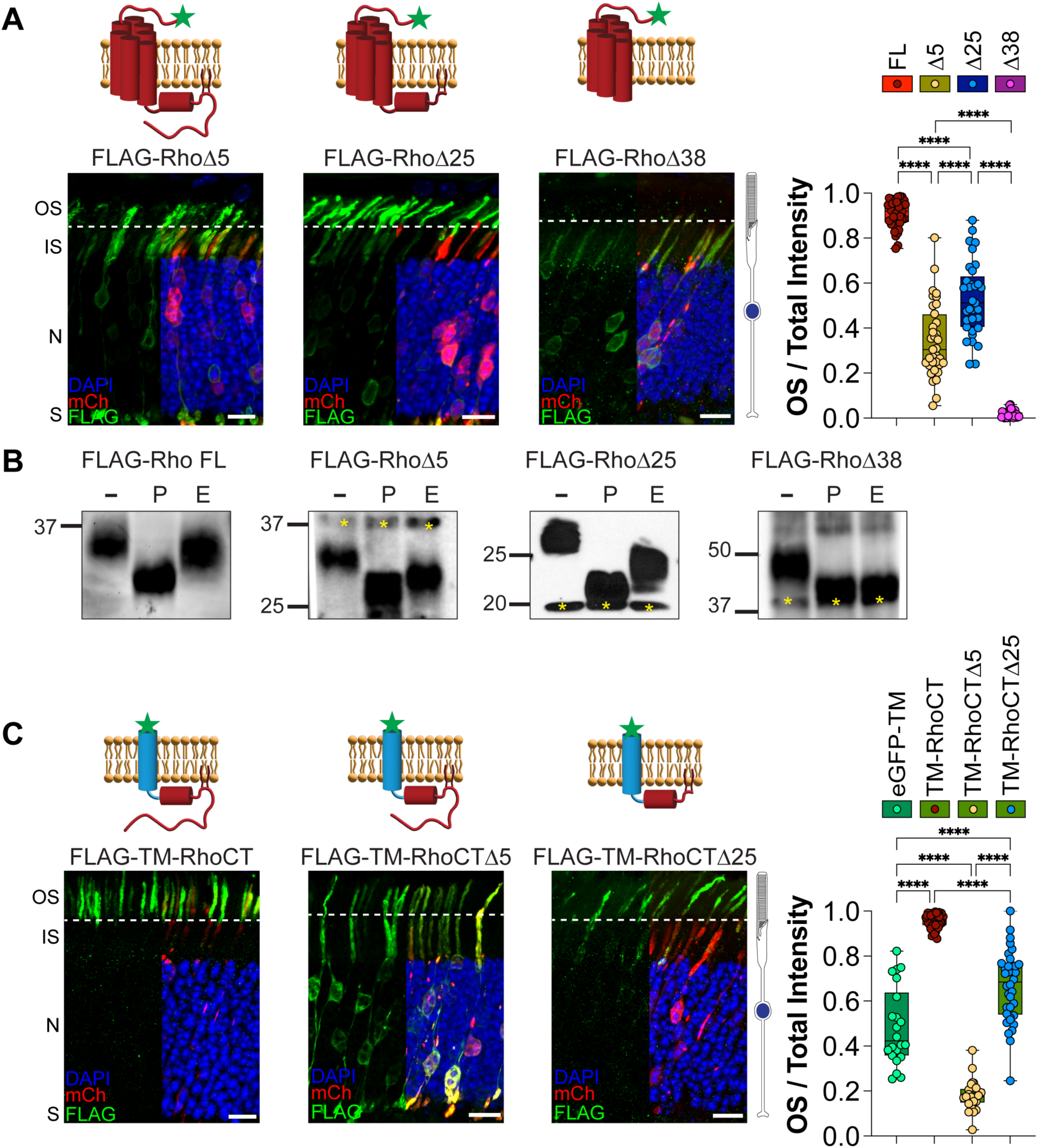
Rhodopsin C-terminal truncations reveal unique localization profiles in WT mouse rods. **A**. The following FLAG-tagged rhodopsin constructs were electroporated into WT mouse rods: rhodopsin lacking its final 5 amino acids (RhoΔ5), 25 amino acids (RhoΔ25) or 38 amino acids (RhoΔ38). Δ5 n=9, 35 images; Δ25 n=10, 33 images; Δ38 n=6, 27 images. **B**. FLAG-tagged rhodopsin constructs immunoprecipitated from electroporated RhoKO mouse retinal lysates were treated with PNGase F or Endo H and analyzed by Western blot. Untreated lysates (-), PNGase F (P), Endo H (E). Sensitivity to both PNGase F and Endo H treatments shows increased electrophoretic mobility for FLAG-RhoΔ38 indicative of retention within the ER. Resistance to Endo H treatment indicates ER exit and normal processing through the conventional secretory pathway for FLAG-Rho, FLAG-RhoΔ5 and FLAG-RhoΔ25. **C**. WT mouse rods were electroporated with FLAG-tagged, single-pass transmembrane domain fused to: intact C-terminal cytosolic tail of rhodopsin (TM-RhoCT), or rhodopsin’s C-terminal tail lacking the final 5 amino acids (TM-RhoCTΔ5) or the final 25 amino acids (TM-RhoCTΔ25). TM-RhoCT n=4, 41 images; TM-RhoCTΔ5 n=6, 24 images; TM-RhoCTΔ25 n=5, 36 images. Bar graphs show the quotient between the outer segment signal over the total signal for each construct. FLAG staining (green), mCherry (red) labels transfected rod cells, and DAPI (blue) used to counter-stain nuclei. Scale Bar, 10 µm.

It is known that rhodopsin utilizes a conventional secretory pathway through the ER and Golgi (2, 30, 41). We performed deglycosylation experiments to determine whether the trafficking route of our rhodopsin truncations is altered. FLAG-Rho mutants were expressed in RhoKO rods to prevent endogenous rhodopsin from influencing the trafficking pathway. FLAG-Rho constructs were immunoprecipitated and then treated with deglycosylation enzymes before protein mobility was assessed by Western blot. The enzyme Peptide-N-Glycosidase F (PNGase F) cleaves any type of N-linked glycans, and results in an increased mobility shift of Rho-FL compared to untreated control. Endoglycosidase H (Endo H) is an enzyme that cleaves unmodified high-mannose N-linked glycans that are added to proteins in the ER. If the protein reaches the Golgi complex, its N-linked glycan moieties are modified and no longer sensitive to Endo H cleavage. Rho-FL is resistant to Endo H confirming that it traffics through the conventional ER-to-Golgi route (**Figure 2B**), equivalent to endogenous rhodopsin (41). We next tested RhoΔ5 and RhoΔ25 and found they were both sensitive to PNGase F but resistant to Endo H suggesting they use a conventional secretory pathway through the Golgi complex (**Figure 2B**). In contrast, we found that RhoΔ38 was sensitive to both PNGase F and Endo H treatment suggesting that it does not enter the Golgi complex (**Figure 2B**). Together with our localization data showing that RhoΔ38 cannot reach the outer segment in WT rods, we establish that RhoΔ38 does not leave through an unconventional pathway but is retained in the ER.

Rhodopsin mutations have previously been shown to result in ER retention when heterologously expressed in cell culture (42, 43). It was further shown that mutant rhodopsin can be ubiquitinated and degraded in both cell culture and mouse models (44). Therefore, we expressed our rhodopsin truncation mutants in cell culture and assessed their localization pattern and rate of degradation. FLAG-tagged rhodopsin constructs were transiently transfected in AD-293 cells and their localization was initially analyzed by staining fixed cells with both a plasma membrane marker, Na/K ATPase, and ER marker, calnexin (**Sup Figure 2**). We found FLAG-Rho FL and FLAG-RhoΔ5 were primarily localized to the plasma membrane where they co-localized with Na/K-ATPase. FLAG-RhoΔ25 appeared to have a varied localization pattern between the ER and the plasma membrane suggesting that RhoΔ25 is not processed normally in AD-293 cells. We observed FLAG-RhoΔ38 was restricted to the ER, similar to its localization in rod photoreceptors. We then tested protein stability by performing cycloheximide (CHX) chase experiments. CHX inhibits translation, preventing the addition of newly synthesized proteins. By taking time points after CHX addition, we can determine the half-life of an expressed protein. Unstable proteins or proteins that become targets for degradation will show a faster reduction in protein levels over time. We did not detect any significant difference in the rate of protein degradation when comparing FLAG-Rho FL to any of the truncated rhodopsin constructs (**Sup Figure 2**). Together, these results suggest that truncated rhodopsin constructs are stable even when retained in the ER.

The 7-TM core structure of rhodopsin is known to provide structural integrity and dimerization interface, so we wanted to assess the information encoded within each of the C-terminal rhodopsin truncations in isolation. We chose to append rhodopsin’s C-terminus to a FLAG-tagged, inert, single-pass transmembrane domain from an activin receptor, previously shown to have no specific targeting signal by itself (34). We expressed the complete 38 amino acid cytosolic C-terminus (FLAG-TM-RhoCT) and found it behaves like full-length rhodopsin, showing an outer segment targeted profile (mean OS/Total Intensity value 0.957±0.033, **Figure 2C**). This is consistent with previously published results using the activin TM domain fused to GFP and rhodopsin C-terminus (34). Removing the QVAPA outer segment targeting signal (FLAG-TM-RhoCTΔ5) resulted in a significant mislocalization from the outer segment to the cell body (mean OS/Total Intensity value 0.181±0.069, **Figure 2C**). This data further supports the idea that a C-terminal mistargeting signal becomes exposed when QVAPA is removed. Further truncation of rhodopsin’s C-terminus (FLAG-TM-RhoCTΔ25) increased outer segment localization, but not to full-length values, suggesting an untargeted localization profile (mean OS/Total Intensity value 0.664±0.157, **Figure 2C**). Together our data shows that the targeting information encoded within rhodopsin’s C-terminus is conserved whether it is in the context of the core structure of rhodopsin or appended onto an inert transmembrane domain.

### Rhodopsin’s QVAPA targeting signal can alleviate ER retention and restore localization of an untargeted rhodopsin to the outer segment

We found that RhoΔ25 behaves as an untargeted protein with the same localization profile as an untargeted eGFP-TM construct. It was previously shown that appending the C-terminal 8 residues of rhodopsin to untargeted constructs could drive specific outer segment localization (25). We appended the QVAPA targeting motif to the RhoΔ25 construct to test whether we could restore outer segment targeting (FLAG-RhoΔ25+QVAPA). In WT rods, the addition of the QVAPA sequence specifically targeted RhoΔ25 to the outer segment so it resembled full-length rhodopsin (OS/Total Intensity value of 0.861±0.082, **Figure 3A**).

**Figure 3:**
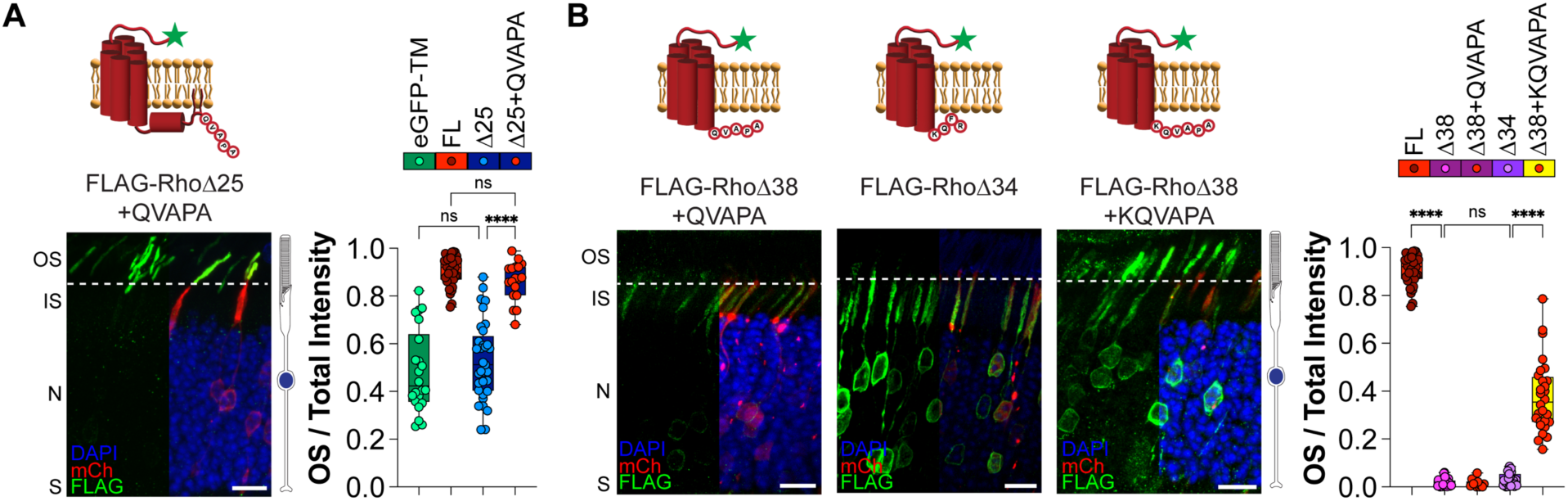
Rhodopsin’s QVAPA outer segment targeting signal requires transmembrane stabilization to alleviate ER retention. **A**. WT mouse rods electroporated with FLAG-tagged rhodopsin lacking the final 25 amino acids fused to QVAPA. Δ25+QVAPA n=4, 18 images. **B**. WT mouse rods electroporated with the following FLAG-tagged rhodopsin constructs: rhodopsin lacking the final 34 amino acids (RhoΔ34) or lacking the final 38 amino acids fused with either QVAPA (RhoΔ38+QVAPA) or KQVAPA (RhoΔ38+KQVAPA). Δ34 n=6, 31 images; Δ38+QVAPA n=7, 32 images; Δ38+KQVAPA n=4, 28 images. Bar graphs show the quotient between the outer segment signal over the total signal for each construct. FLAG staining (green), mCherry (red) labels transfected rod cells, and DAPI (blue) used to counter-stain nuclei. Scale Bar, 10 µm.

We next wanted to determine whether we could induce normal ER exit of RhoΔ38 by appending the QVAPA targeting sequence to its C-terminus (FLAG-RhoΔ38+QVAPA). In WT rods, FLAG-RhoΔ38+QVAPA was present in internal membranes surrounding the cell body with no localization to the outer segments, similar to FLAG-RhoΔ38 (OS/Total Intensity value of 0.015±0.018, **Figure 3B**). Since FLAG-RhoΔ38 lacks the positive lysine residue at the end of the seventh transmembrane helix, the protein topology is likely disrupted leading to the misfolded phenotype. To circumvent misfolding due to transmembrane instability, we extended the rhodopsin C-terminus to FLAG-RhoΔ34 providing the necessary lysine residue to ensure the seventh transmembrane helix is correctly embedded. We found that FLAG-RhoΔ34 is still retained in the ER with an OS/Total Intensity value of 0.029±0.027 (**Figure 3B**). When we appended the targeting signal including the necessary lysine residue onto RhoΔ38 (FLAG-RhoΔ38+KQVAPA), we found it was localized in the outer segment and around the cell body. The localization profile value was significantly increased, suggesting the KQVAPA alleviated ER retention of RhoΔ38 (OS/Total Intensity value of 0.378±0.150, **Figure 3B**). Our data suggests that in addition to its classical role as an outer segment targeting signal, the C-terminal QVAPA sequence of rhodopsin can also relieve ER retention if the core helical bundle of rhodopsin is maintained.

### Examining the intrinsic outer segment elongation properties of rhodopsin C-terminus truncations

RhoKO rods develop a small, rudimentary outer segment compartment filled with membrane tubules. These residual membranes are densely packed with the other outer segment resident proteins, whose trafficking is largely unaffected by the loss of rhodopsin (31). When we express our untargeted membrane reporter, eGFP-TM, into RhoKO rods we find enrichment in the rudimentary outer segment with a large fraction residing within the cell body (OS/Total Intensity value of 0.231±0.115, **Figure 4A**). In contrast, FLAG-Rho FL is specifically localized to the rudimentary outer segments, which appear elongated relative to eGFP-TM (OS/Total Intensity value of 0.739±0.095, **Figure 4A**). We next tested the rhodopsin truncations and found that all of them were largely absent from the rudimentary outer segments and failed to induce elongation of this compartment (OS/Total Intensity values: Δ5 0.089±0.110; Δ25 0.050±0.038; Δ38 0.011±0.014; **Figure 4A**). This suggests that the entire C-terminus of rhodopsin is required for normal elongation of the outer segment.

**Figure 4:**
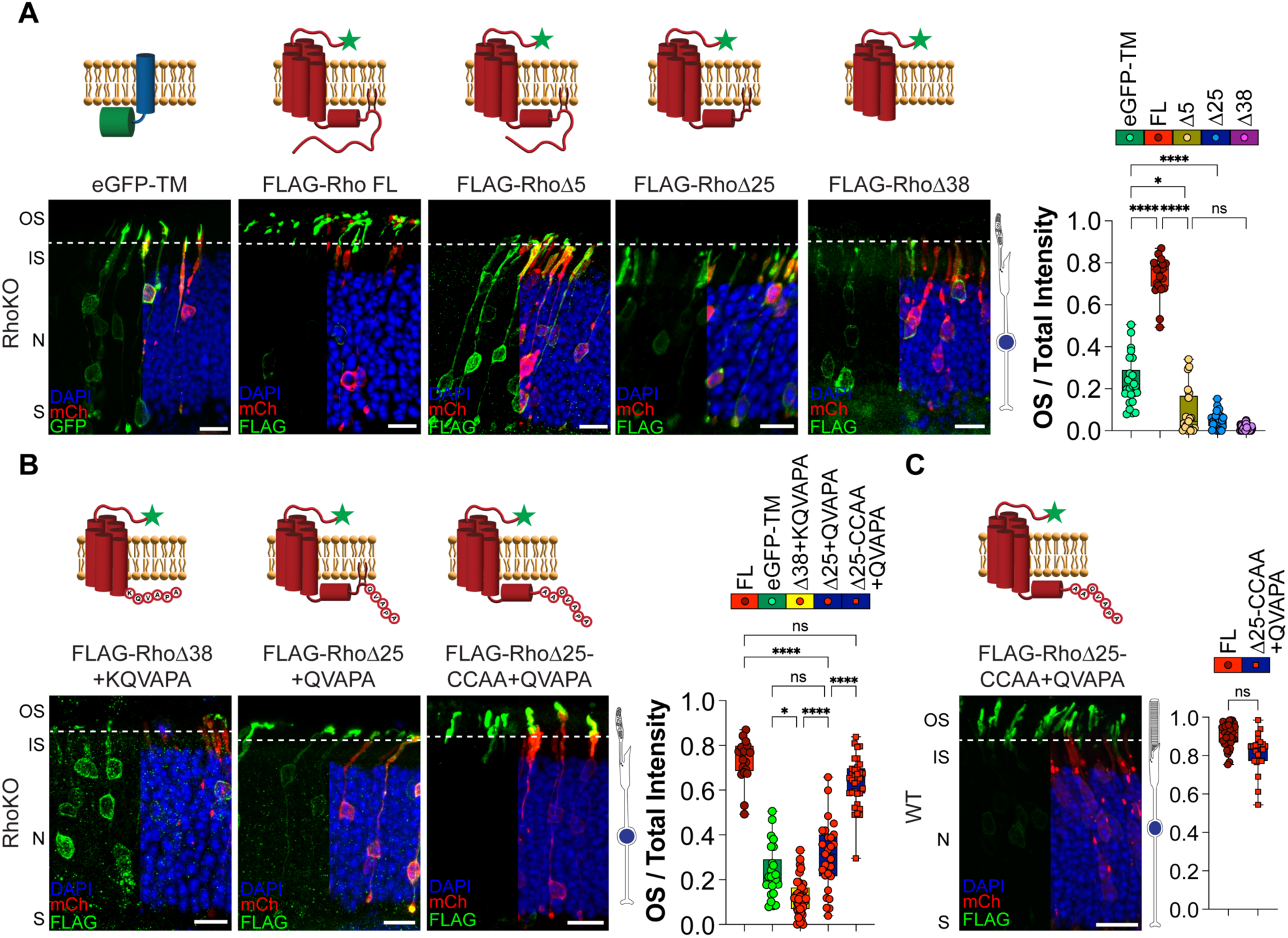
Rhodopsin’s helix-8, but not cysteine anchors, is required for proper outer segment elongation in rhodopsin knockout rods. **A**. RhoKO mouse rods were electroporated with an eGFP-tagged transmembrane segment 1 from mGluR1 (eGFP-TM) construct or the following FLAG-tagged rhodopsin constructs: full-length rhodopsin (Rho FL), rhodopsin lacking its final 5 amino acids (RhoΔ5), 25 amino acids (RhoΔ25) or 38 amino acids (RhoΔ38). eGFP-TM n=6, 26 images; FL n=3, 21 images; Δ5 n=7, 21 images; Δ25 n=8, 27 images; Δ38 n=5, 40 images. **B**. RhoKO rods were electroporated with FLAG-RhoΔ38+KQVAPA, FLAG-RhoΔ25+QVAPA or FLAG-RhoΔ25-CCAA+QVAPA. Δ38+KQVAPA n=6, 39 images; Δ25+QVAPA n=4, 27 images; Δ25-CCAA+QVAPA n=6, 33 images. **C**. Image showing WT rods electroporated with FLAG-RhoΔ25-CCAA+QVAPA (n=4, 21 images). Bar graphs show the quotient between the outer segment signal over the total signal for each construct. FLAG staining or GFP (green), mCherry (red) labels transfected rod cells, and DAPI (blue) used to counter-stained nuclei. Scale Bar, 10 µm. To the right is a cartoon of a RhoKO rod photoreceptor, showing disrupted outer segment structure.

We wanted to determine if driving additional delivery of rhodopsin’s core structure into the rudimentary outer segment of RhoKO mice could elongate this compartment as we observed with FLAG-Rho FL. We previously found that QVAPA restores outer segment localization of rhodopsin truncated constructs in WT mice (**Figure 3**), so we electroporated FLAG-RhoΔ38+KQVAPA or FLAG-RhoΔ25+QVAPA into RhoKO rods. We found that FLAG-RhoΔ38+KQVAPA was primarily localized around the cell body failing to elongate the rudimentary outer segments (OS/Total Intensity value of 0.122±0.079, **Figure 4B**). The FLAG-RhoΔ25+QVAPA construct showed more robust localization within the rudimentary outer segments; however, it was unable to extend the compartment and was not significantly different from the eGFP-TM untargeted membrane construct (OS/Total Intensity value of 0.304±0.155, **Figure 4B**). This result was surprising as we previously found that FLAG-RhoΔ25+QVAPA was properly targeted to the outer segment in WT rods and expected that restoring outer segment delivery of the core structure of rhodopsin would be able to elongate the rudimentary outer segments similar to FLAG-Rho FL.

We postulated that perhaps the presentation of the QVAPA targeting motif, while fully targeting RhoΔ25 in WT rods where endogenous rhodopsin is present, could be constrained when expressed in isolation in RhoKO rods. We wanted to try to destabilize helix-8 to allow for a better presentation of the QVAPA motif, so we first tested several helix-8 mutations in the context of RhoΔ25 to ensure these constructs had proper ER exit (**Sup Figure 3**). We found that mutating the cysteine residues at positions 322 and 323 to alanine did not affect the overall localization of FLAG-RhoΔ25 in WT and RhoKO rods (**Sup Figure 3**). These cysteine residues are palmitoylated to help provide membrane anchoring of helix-8, so we tested whether the absence of membrane stabilization could relieve potential steric hindrances of the QVAPA targeting motif. We found that electroporation of FLAG-RhoΔ25-CCAA+QVAPA in RhoKO rods resulted in robust outer segment delivery and elongation of rudimentary outer segments, comparable to FLAG-Rho FL (OS/Total Intensity value of 0.636±0.111, **Figure 4B**). Additionally, we confirmed that FLAG-RhoΔ25-CCAA+QVAPA in WT rods was localized to the outer segment, similar to Rho FL (OS/Total Intensity value of 0.815±0.102, **Figure 4C**). From these results, we conclude that targeted delivery of the core structure of rhodopsin with helix-8 is sufficient for outer segment elongation in RhoKO rods.

### Rhodopsin mutants with partial mislocalization cannot elongate outer segments in RhoKO rods

Our data suggests that the localization profile of a particular rhodopsin construct in WT mice does not predict whether that same construct can elongate rudimentary outer segments in RhoKO rods. To further investigate this phenomenon, we tested rhodopsin mutations previously shown to have no effect on rhodopsin localization or have partial mislocalization from the outer segment to the inner segment. A previous publication found that a palmitoylation defective rhodopsin mutant resulted in normal rhodopsin localization and outer segment formation (45). We electroporated a FLAG-Rho-CCAA mutant into WT mice and found it to be properly localized to the outer segment (OS/Total Intensity value of 0.935±0.039, **Figure 5A,D**). When FLAG-Rho-CCAA was expressed in RhoKO rods, it localized and elongated the rudimentary outer segments similar to FLAG-Rho FL (OS/Total Intensity value of 0.715±0.169, **Figure 5A,D**). This result confirms the reliability of our electroporation assay and that rhodopsin palmitoylation is not required for localization or outer segment elongation, as was previously shown.

**Figure 5:**
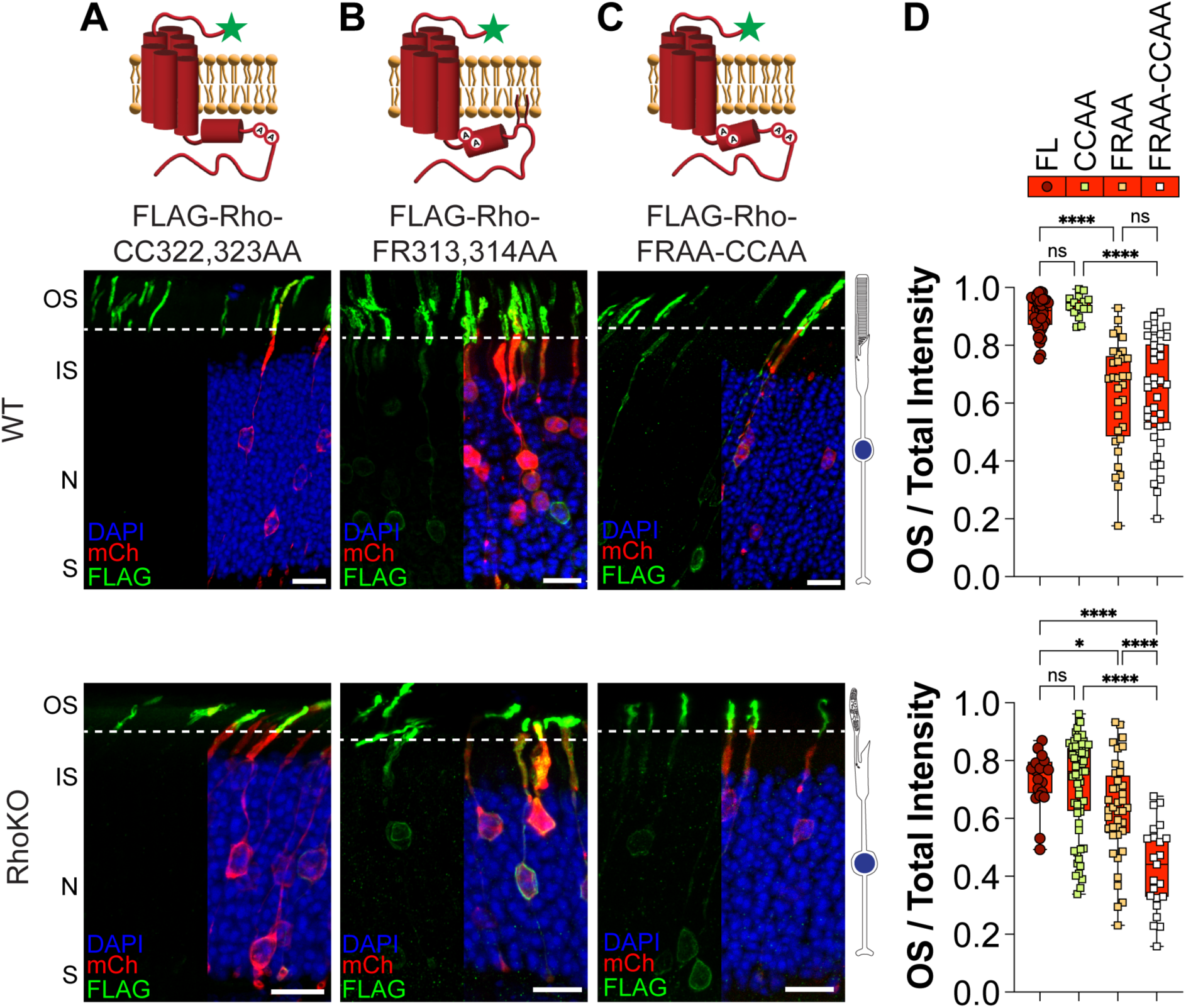
Mislocalized full-length rhodopsin mutants in WT rods are unable to properly build outer segments in RhoKO rods. **A**. FLAG-tagged rhodopsin with a CC(322,323)AA mutation in helix-8, disrupting the palmitoylated cysteine membrane anchor, shows outer segment targeting comparable to full-length rhodopsin in WT rods (top panel). In RhoKO rods, the CCAA mutant elongates the rudimentary outer segment like FL (bottom panel). **B**. FLAG-tagged rhodopsin with a FR(313,314)AA mutation in helix-8 displays a mislocalization profile in WT rods (top panel); and is unable to sufficiently elongate the outer segment in RhoKO rods (bottom panel). **C**. The double mutant combining FRAA and CCAA mutations phenocopies the FRAA mislocalization profile in WT rods (top panel) and runty outer segment profile in RhoKO rods (bottom panel). **D**. Bar graphs show the quotient between the outer segment signal over the total signal for each construct described in **A-C**. WT analysis: CCAA n=4, 15 images; FRAA n=8, 32 images; FRAA-CCAA n=5, 38 images. RhoKO analysis: CCAA n=9, 61 images; FRAA n=8, 40 images; FRAA-CCAA n=6, 23 images. FLAG staining (green), mCherry (red) labels transfected rod cells, and DAPI (blue) used to counter-stained nuclei. Scale Bar, 10 µm.

The F313 and R314 residues in Helix-8 are highly conserved in GPCRs localized to the cilium (46, 47). F313 was shown to form hydrophobic interactions with L68 of TM1 and mutating this phenylalanine to arginine resulted in partial mislocalization of rhodopsin to the inner segment (19). We electroporated a FLAG-Rho-FRAA mutant construct in WT rods and confirmed that destabilizing the interhelix hydrophobic interaction resulted in a fraction of rhodopsin mislocalized to the cell body (OS/Total Intensity value of 0.630±0.187, **Figure 5B,D**). We then expressed the FLAG-Rho-FRAA construct in RhoKO rods and found that despite strong localization in rudimentary outer segments that appeared larger, a fraction remained localized within the cell body (**Figure 5B**). This resulted in FLAG-Rho-FRAA having an OS/Total Intensity value significantly lower than FLAG-Rho FL (OS/Total Intensity value of 0.630±0.169, **Figure 5D**). Together, our data suggest that destabilizing interhelical hydrophobic interactions prevents sufficient rhodopsin localization to the outer segment, likely failing to properly form this compartment.

We also tested a rhodopsin construct that combined the FRAA and CCAA mutations and found that in WT it behaved similarly to the FRAA with partial mislocalization from the outer segment (OS/Total Intensity value of 0.630±0.192, **Figure 5C,D**). However, when expressed in RhoKO rods we found a synergistic effect with increased mislocalization and reduced elongation compared to the FRAA (OS/Total Intensity value of 0.430±0.148, **Figure 5C,D**). This data suggests that precise outer segment targeting of rhodopsin is a critical driver for normal outer segment formation.

## DISCUSSION

Our study provides a comprehensive analysis of the localization profile for rhodopsin C-terminus mutants expressed in either WT or RhoKO rods. Analyzing rhodopsin constructs *in vivo* is important as cell culture experiments do not provide the native environment or trafficking demands unique to rod photoreceptors. In our data, we can determine whether the expressed rhodopsin construct is localized to the outer segment in the presence or absence of endogenous rhodopsin and whether the construct can elongate the rudimentary outer segment in RhoKO rods. Outer segment targeting and building efficacy of each rhodopsin mutant was determined by comparing it to either a full-length rhodopsin or an untargeted eGFP-TM construct. The initial RhoΔ5, RhoΔ25 and RhoΔ38 truncation constructs we tested helped to establish distinct profiles: mistargeted, untargeted, and ER retained, respectively (**Figure 6**). Despite mistargeted and untargeted constructs accumulating in the outer segment of WT rods, we found these constructs were unable to extend the rudimentary outer segments in RhoKO rods (**Figure 2A, 4A**). In particular, we were surprised that the untargeted RhoΔ25 construct with strong outer segment localization in WT rods was not even enriched in the rudimentary outer segment of RhoKO rods. Contrasting the untargeted eGFP-TM construct, which accumulated in the outer segments of WT and RhoKO rods. The possibility of RhoΔ25 forming heterodimers with endogenous rhodopsin could explain the nuance observed in this result but was not examined in this study.

**Figure 6:**
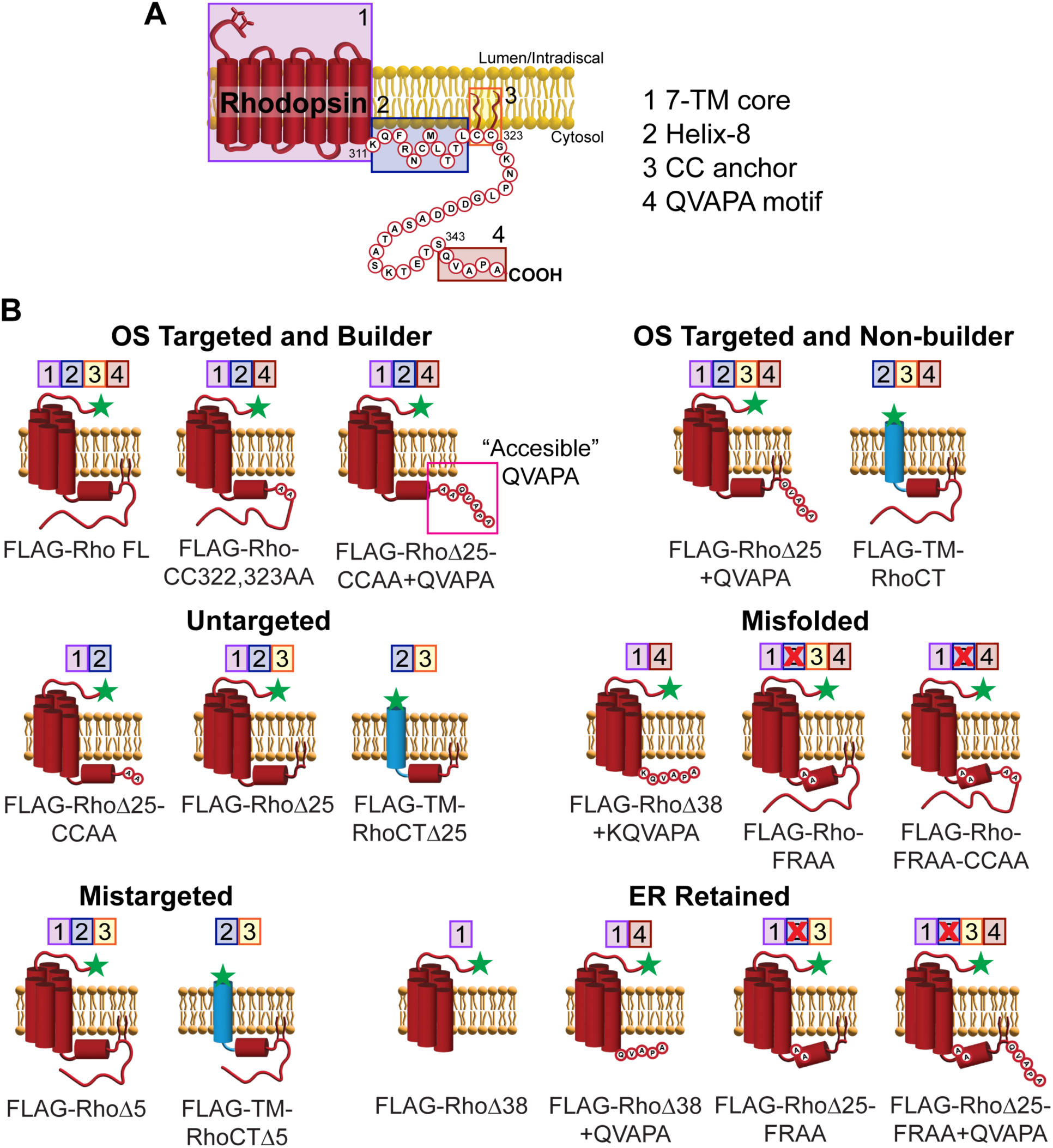
Quick-glance summary of rhodopsin constructs localization pattern. **A**. Diagram of mouse rhodopsin highlighting key topological features analyzed in this study: (1) rhodopsin 7-TM core, (2) amphipathic helix-8, (3) CC anchor and (4) QVAPA motif. **B**. FLAG-tagged rhodopsin constructs tested in this study are grouped according to their profiles. Localization-specific classifications were derived from the profiles for Rho FL (OS Targeted), eGFP-TM (Untargeted), RhoΔ5 (Mistargeted) and eGFP-Cb5-TM (ER Retained). Builder designation is determined by the capacity to extend the small outer segment in RhoKO rods. The Misfolded group was designated as constructs that contain the QVAPA targeting motif but are mislocalized from the outer segment in WT rods. Topological features present in each construct are listed above the cartoon. Results are also summarized in **Supplemental Table 1.**

The disconnect between a construct’s localization pattern in WT and its outer segment elongation efficiency in RhoKO was also apparent when we began appending the QVAPA motif onto truncated constructs. By appending the QVAPA targeting signal to RhoΔ25, we fully rescued its localization to the outer segment in WT rods but not its building efficacy in RhoKO rods (**Figure 3A, 4B**). Only when we disrupted the CC anchor in the RhoΔ25+QVAPA construct were we able to rescue outer segment elongation in RhoKO rods (**Figure 4B**). Membrane attachment provided by the CC anchor helps position helix-8 adjacent to the membrane, which we postulate could prevent access to the QVAPA motif.

When we tested an alternative helix-8 disrupting mutation, F313 and R314 to alanines, in the RhoΔ25+QVAPA construct it resulted in retention in the ER in both WT and RhoKO rods (**Sup Figure 3**). This suggests that an ER retention supersedes targeting, which we also observed to be true when we appended QVAPA onto RhoΔ38, another ER retained mutant (**Figure 3B**). Only when a locking lysine was added before the QVAPA targeting motif (RhoΔ38+KQVAPA) could we restore outer segment localization in WT rods (**Figure 3B**). However, the presence of the same locking lysine when in the absence of QVAPA resulted in an ER retention phenotype (**Figure 3B**), which suggests that QVAPA plays a role in supporting ER exit.

Previous studies of ciliary GPCRs have found that the intracellular loop 3 encodes ciliary targeting (48, 49). By examining the C-terminus of rhodopsin in isolation we found similar localization patterns as with rhodopsin’s 7-TM core suggesting that any additional signals within the core structure of rhodopsin are negligible.

Previous work analyzed C-terminally truncated human rhodopsin expressed in frog rods (20). While our results from mouse rods generally align with what was observed for frog rods, one notable difference is for rhodopsin truncations beyond the CC anchor. In frogs, severe truncations of human rhodopsin (RhoΔ27, RhoΔ32, RhoΔ38) resulted in predominant localization to the outer segment (20). In this study, we find mouse rhodopsin truncations (RhoΔ34 and RhoΔ38) to be retained in the ER. A key difference between our studies is that the previous study fused Dendra2 to the C-terminus of truncated rhodopsin, whereas we appended a FLAG-tag to the N-terminus. One possibility is that the Dendra2 may have provided additional protein stability to the truncated human rhodopsin allowing for ER exit. Another possible explanation could be that frog rods process rhodopsin, or transmembrane proteins in general, differently than mouse rods. This scenario could be likely due to the larger volume of frog outer segments predicted to result in higher membrane trafficking demand compared to mouse rods (34). Regardless of the conflicting results, our study finds even if RhoΔ38 was able to reach the outer segment in WT rods, shown by appending KQVAPA, it would likely be unable to generate a proper outer segment (**Figure 3B, 4B**). Thus, our system provides further context for rhodopsin mutants by determining whether they are sufficient to elongate outer segments in RhoKO rods. One limitation to our study is that we don’t examine whether rhodopsin constructs are properly building disc membranes within the outer segment. This would require ultrastructural analysis, which is beyond the scope of this study.

## MATERIALS AND METHODS

### Animals

Albino CD-1 wild-type mice were ordered from Charles River Laboratories (Strain # 022; Mattawan, MI). Rhodopsin knockout (RhoKO) mice were a generous gift from Vadim Arshavsky (Duke University)(37). Mice were handled following protocols approved by the Institutional Animal Care and Use Committees at the University of Michigan (registry number A3114-01). As there are no known gender-specific differences related to vertebrate photoreceptor development and/or function, male and female mice were randomly allocated to experimental groups. All mice were housed in a 12/12-hour light/dark cycle with free access to food and water.

### *In vivo* electroporation of mouse retinas

DNA constructs were electroporated into the retinas of neonatal mice using the protocol as originally described in (50), but with modifications as detailed in (51). Briefly, P0-P2 mice were anesthetized on ice, had their eyelid and sclera punctured at the periphery of the eye with a 30- gauge needle, and were injected with 0.25 - 0.5 µL of concentrated plasmid in the subretinal space using a blunt-end 32-gauge Hamilton syringe. A plasmid mixture consists of 2 µg/µL of gene of interest and 1 µg/µL of rhodopsin promoter driving expression of mCherry (pRho:mCherry) as an electroporation marker to identify transfected cells. A complete list of constructs can be found in **Supplemental Table 1**. The positive side of a tweezer-type electrode (BTX, Holliston, MA) was placed over the injected eye and five 100 V pulses of 50 milliseconds were applied using an ECM830 square pulse generator (BTX). Neonates were returned to their mother until collection at P20-P22.

### Cell culture

All cell culture experiments were performed using AD-293 cells (Agilent Technologies, 240085; RRID:CVCL_9804). Cells were either plated onto poly-L-lysine glass coverslips (Corning, 354085; Glendale, AZ) in a 24-well plate for imaging or in a 10 cm plate for biochemical assays and grown overnight at 37°C with 5% CO_2_. Transient transfections were performed 1 day after seeding and experimental collection was performed 2 days after transfection. 0.5 μg/mL of plasmid DNA was used for transfections by incubating the DNA with 1 μg/mL polyethyleneimine (Sigma, 408727) in serum-free media for 10 min before being added dropwise to cells that had been changed to serum-free media 1 hr prior.

### DNA constructs

A complete list of primers used for cloning can be found in **Supplemental Table 2**. For rod-specific expression, cDNA sequences were cloned between 5’ *AgeI* and 3’ *NotI* cloning sites into a vector driven by a 2.2 kb bovine rhodopsin promoter originally cloned from pRho:DsRed, (gift from Dr. Connie Cepko; Addgene plasmid #11156; http://n2t.net/addgene:11156). For expression in cell culture, inserts from the pRho plasmids were transferred by molecular cloning with AgeI and NotI cloning sites into a pEGFP-N1 (Clontech) plasmid containing the CMV promoter.

### Antibodies

The following commercial primary antibodies were used: mouse monoclonal M2 anti-FLAG (F1804, Sigma-Aldrich); rabbit polyclonal anti-FLAG (F7425, Sigma-Aldrich); goat polyclonal anti-FLAG (ab1257, Abcam); rabbit polyclonal anti-FLAG (PA1-948B, Thermo Fisher); goat polyclonal anti-mCherry (AB0040, OriGene Technologies); rabbit polyclonal anti-Calnexin (C4731, Sigma); and mouse monoclonal anti-Na/K-ATPase (SC58628, Santa Cruz). The following commercial secondary antibodies were used: Donkey anti-mouse AF488 (A21202, Fisher Scientific); Donkey anti-rabbit AF488 (A21206, Fisher Scientific); Donkey anti-sheep (A11015, Fisher Scientific); Donkey anti-sheep AF568 (A21099, Fisher Scientific); Donkey anti-mouse AF647 (A31571, Fisher Scientific); Donkey anti-Sheep AF647 (A21448, Fisher Scientific); and Donkey anti-Rabbit Peroxidase AffiniPure (711-035-152, Jackson ImmunoResearch).

### Immunofluorescence

Electroporated mouse eyes were enucleated and drop-fixed in 4% paraformaldehyde in 1x PBS (AAJ75889K8, Fisher Scientific) at room temperature for 1-2 hours before washing with 1x PBS. Eyecups were dissected and embedded in 4% low-gelling temperature agarose in 1x PBS (A9045; Sigma-Aldrich) and cut into 100 µm thick sagittal sections with a vibratome (Leica Biosystems). Free-floating sections were blocked in 5% donkey serum (NC0629457, Thermo Fisher) and 0.5% Triton X-100 (BP151, Fisher Scientific) in 1x PBS for 1 hour at room temperature. Then incubated with primary antibodies overnight at 4°C, washed with 1x PBS, and then incubated in blocking buffer for 1 hour at room temperature with secondary antibodies along with 10 µg/ml DAPI (102362760, Sigma-Aldrich). Sections were washed in 1x PBS and mounted on slides with ProLong Glass Mounting Media (P36980, Thermo Fisher) and #1.5 coverslips (72204-10, Electron Microscopy Sciences).

Transfected cells on coverslips were fixed in 4% paraformaldehyde for 1 hour. Cells were then washed with 1x PBS, permeabilized for 5 minutes with 0.5% Triton X-100 in 1x PBS, washed with 1x PBS and nonspecific binding was blocked with a 1-hour incubation in 1x PBS containing 0.5% Triton X-100 and 5% normal donkey serum. Coverslips were then incubated overnight at 4°C with primary antibody diluted in blocking buffer (antibody information listed above), washed in 1x PBS, and incubated for 1-2 hours at 22°C with 10 μg/mL DAPI and donkey secondary antibodies conjugated with Alexa Fluor 488, 568, or 647. Coverslips were washed in 1x PBS and mounted on slides with Immunomount for imaging.

Images were acquired using a Zeiss Observer 7 inverted microscope equipped with a 63x oil-immersion objective (1.40 NA), LSM 800 confocal scanhead outfitted with an Airyscan superresolution detector controlled by Zen 5.0 software (Carl Zeiss Microscopy; White Plains, NY). Manipulation of images was limited to adjusting the brightness level, image size, rotation and cropping using FIJI (ImageJ, https://imagej.net/Fiji; ImageJ2 Version 2.14.0/1.54f; Build c89e8500e4).

### Image Quantification

All imaging processing and analysis was performed using the FIJI imaging software. A max intensity projection image was produced from a Z-stack image to quantify the ratio of the outer segment (OS) and total fluorescence intensity of the electroporated rod photoreceptors. The max stack image was subjected to a 20-pixel rolling-ball background subtraction and threshold set using the Triangle algorithm. When needed, images were subjected to additional manual thresholding to exclude background fluorescence. Binary masks were then produced from the threshold image and fluorescence areas assigned as regions-of-interest (ROI). ROIs were then curated to exclude non-specific staining (e.g. secondary antibody binding to blood vessels or immune cells) and generate the total raw integrated density value for a given image. Using the soluble mCherry transfection marker as a guide, an additional “cell body” ROI was added to select the region that contains the inner segment, outer nuclear layer, and synaptic terminals. To calculate the outer segment “OS” value, the raw integrated density of “cell body” ROI was subtracted from the total ROI. For each image, the OS targeting ratio was determined by the resulting quotient values from a simple “OS/total” calculation, where 1.0 is fully targeted and 0.0 fully excluded from the outer segment.

### Deglycosylation Assay of FLAG-Rhodopsin constructs

Eyecups from electroporated *Rho^-/-^* mice were collected at P21, screened for positively transfected regions and pooled into one tube. A minimum of 6 transfected eyecups were used for each immunoprecipitation/deglycosylation assay. Eyecups were lysed in 1 mL 1% DDM in PBS with protease inhibitors. Lysates were cleared at 15,000 rpm for 10 min at 4°C and supernatant collected for FLAG immunoprecipitation. FLAG-Rho FL n=3; FLAG-RhoΔ5 n=2; FLAG-RhoΔ25 n=3; FLAG-RhoΔ38 n=2.

For PNGase F treatment, lysate is combined with 2 mL each Glycoprotein Denaturing Buffer, GlycoBuffer 2 and 10% NP-40, 1 mL PNGase F (P0708L, New England Biolabs), and H2O as necessary to a final volume of 20 mL. The reaction was incubated for 1 hour at 37°C. For Endo H (P0702L, New England Biolabs) treatment, the same procedure was followed, except that GlycoBuffer 3 was used instead of GlycoBuffer 2. For control experiments, reactions used GlycoBuffer 2 without enzymes. The reaction was terminated by cooling on ice or freezing at - 20°C. 5x sample buffer with 100 mM dithiothreitol (DTT) was added to each sample. Westerns blotting was performed by running lysates on SDS-PAGE using AnykD Criterion TGX Precast Midi Protein Gels (5671124, Bio-Rad) was followed by transfer at 66 mV for 120 minutes onto Immun-Blot Low Fluorescence PVDF Membrane (1620264, Bio-Rad). Membranes were blocked using Intercept Blocking Buffer (927-70003; LI-COR Biosciences, Omaha, NE). Primary antibodies were diluted in 50% / 50% of Intercept / PBST and incubated overnight rotating at 4°C. The next day, membranes were rinsed 3 times with PBST before incubating in the corresponding secondary donkey antibodies conjugated with Alexa Fluor 680 or 800 (LiCor Bioscience) in 50% / 50% / 0.02% of Intercept / PBST / SDS for 2 hours at 4°C. Bands were visualized and quantified using the Odyssey CLx infrared imaging system (LiCor Bioscience). Images of the uncropped Western blots can be found in **Sup Figure 4**.

### Experimental design and statistical analyses

All *in vivo* electroporated mouse retinas were analyzed at postnatal day 20-22. Biological replicates (n’s) were defined as a positively expressing eye, with a minimum of 3 n’s analyzed for every DNA construct. Each “n” consisted of multiple Z-stack images within the expressing retinal tissue. Animals of both sexes were used for all experimental models and were not distinguished in the data. Statistical analysis was performed using Prism 10.0 (GraphPad Prism Software Version 10.3.1). P-values for the OS targeting quotients were determined by using the mean quotient value from each individual “n” (average of averages). Constructs in either WT or RhoKO mice were compared using a 2-way ANOVA (Tukey test to correct for multiple comparisons). All p-values can be found in the multicomparison graphs provided in **Sup Figure 1**.

## ACKNOWLEDGEMENTS

This work was supported by NIH R00 EY025732 (JNP), R01 grant EY032491 (JNP) and NIH T32 grant EY13934 (JYMM). A Career Development Award (JNP) and an Unrestricted Grant to the University of Michigan from Research to Prevent Blindness. The University of Michigan Vision Research Center is also supported by an NIH P30 grant EY007003.

## AUTHOR CONTRIBUTIONS

Conceptualization, JYMM and JNP; Methodology, JYMM and JNP; Investigation, JYMM, SH, AMB, CBS and JNP; Formal Analysis, JYMM, AMB and JNP.; Writing – Original Draft, JYMM and JNP; Writing – Review & Editing, JYMM, SH, CBS and JNP.; Funding Acquisition, JYMM and JNP; Supervision, JNP

## FIGURE LEGENDS

**Supplemental Figure 1: 2-way ANOVA p-Value comparisons grids.** Each construct tested in this study was compared to every other construct expressed in WT or RhoKO rods. Red boxes highlight the p-Values shown in the figures and results. Heat-map legend shows significance values from 2-way ANOVA comparisons.

**Supplemental Figure 2: Rhodopsin truncations expressed in AD293 cells show varied localization patterns with no change to their rates of degradation.** Top, FLAG-tagged rhodopsin constructs transfected into AD293 cells (green). Top panels show Na/K ATPase staining (magenta) to mark the plasma membrane with colocalization in white; bottom panels show Calnexin staining (red) to mark the ER with colocalization in yellow. Scale Bar, 10 µm. Bottom, lysates from AD293 cells transfected with FLAG-tagged rhodopsin constructs were collected before and after 4, 8 or 24hr treatment with cycloheximide (CHX) and analyzed by Western blot. CHX-treatment pauses protein synthesis and enables the analysis of degradation rates over time. Graphic representation of the CHX experiments is shown below. Degradation rates between the tested constructs were not significant.

**Supplemental Figure 3: Disruption of helix-8 in RhoΔ25, but not CC anchors, results in failure to exit the ER.** WT or RhoKO mouse rods (top or bottom panels, respectively) electroporated with FLAG-tagged rhodopsin lacking the final 25 amino acids (RhoΔ25) with the following modifications: CC anchor mutated to alanines (CCAA), FR motif residues mutated to alanines (FRAA), or FRAA mutation with C-terminally appended QVAPA (FRAA+QVAPA). Bar graphs show the quotient between the outer segment signal over the total signal for each construct described. WT analysis: Δ25-CCAA n=4, 18 images; Δ25-FRAA n=6, 28 images; Δ25-FRAA+QVAPA n=3, 15 images. RhoKO analysis: Δ25-CCAA n=4, 19 images; Δ25-FRAA n=4, 16 images; Δ25-FRAA+QVAPA n=6, 16 images. FLAG staining (green), mCherry (red) labels transfected rod cells, and DAPI (blue) used to counter-stained nuclei. Scale Bar, 10 µm.

**Supplemental Figure 4: Uncropped images of Western blots used in Figure 2B**.

**Supplemental Table 1: Summary of all transgenic constructs presented in this study.**

**Supplemental Table 2: Oligonucleotide primers to generate and sequence confirm constructs used in this study.**

