## Supplemental Materials for "Targeted delivery of rhodopsin’s assembled core is required for outer segment extension in mouse rod photoreceptors"

**Supplemental Figure 1**

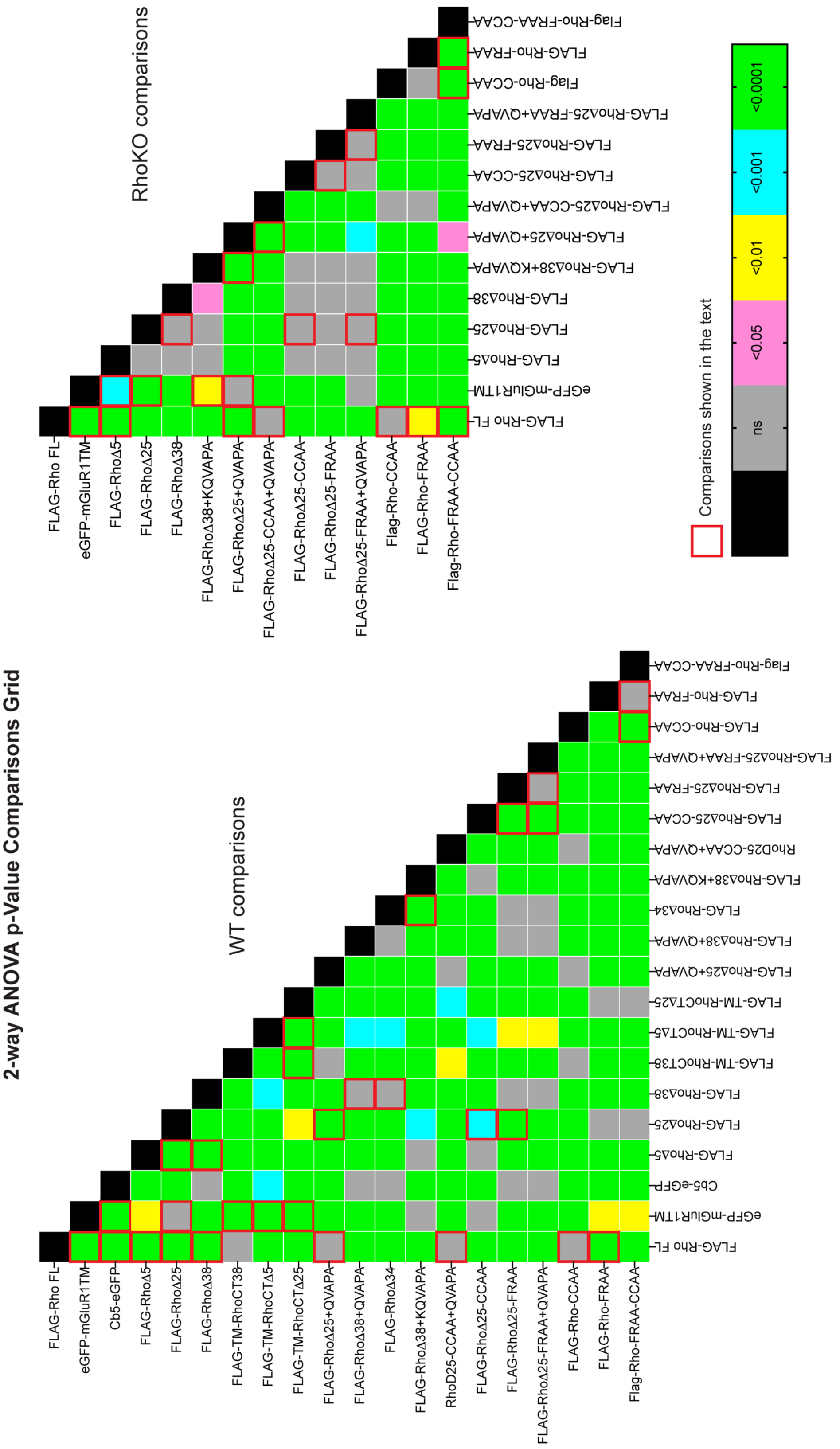

**Supplemental Figure 2**

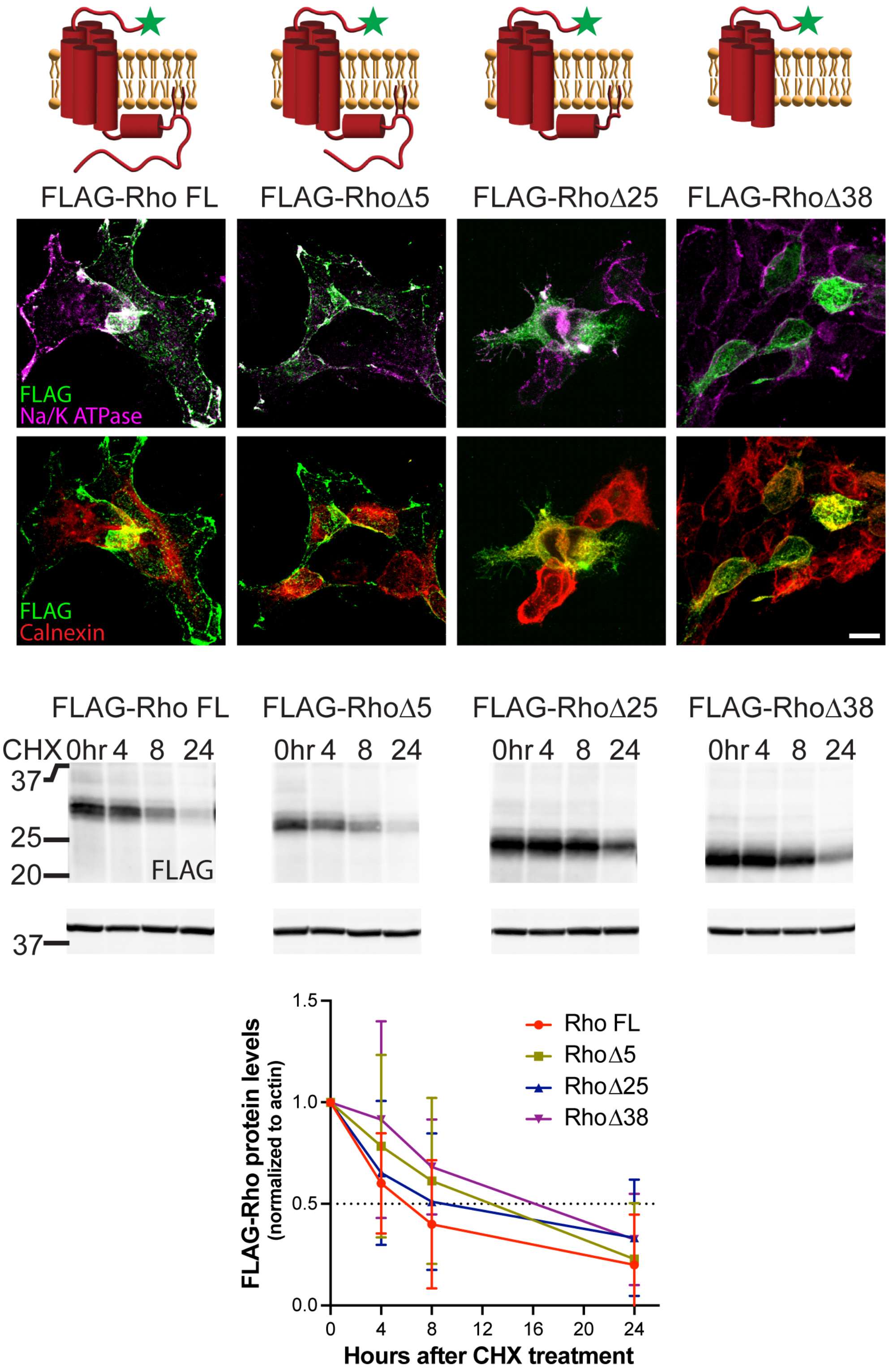

**Supplemental Figure 3**

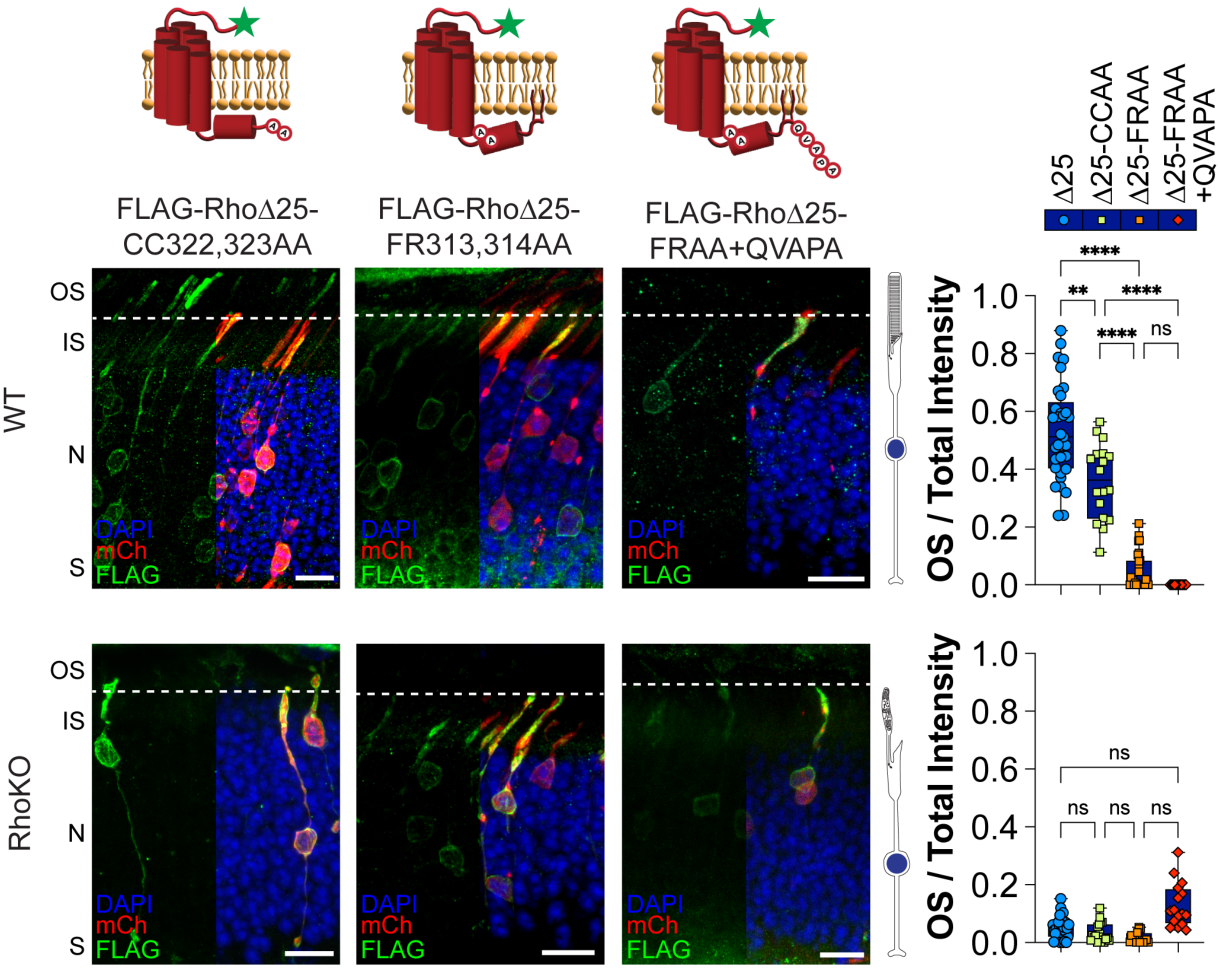

### Supplemental Figure 4

Figure 2B - Rho FL

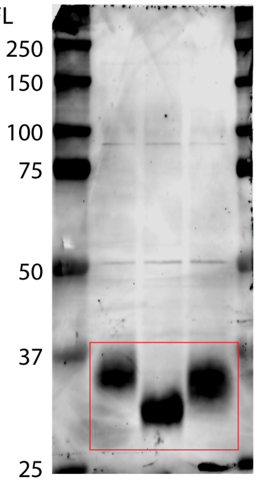


Figure 2B - RhoΔ5

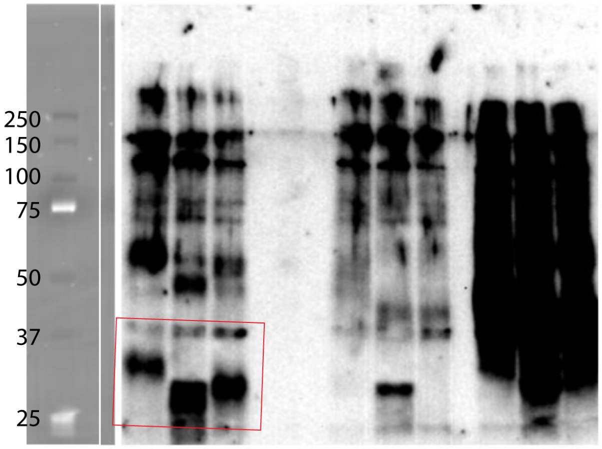


Figure 2B - RhoΔ25

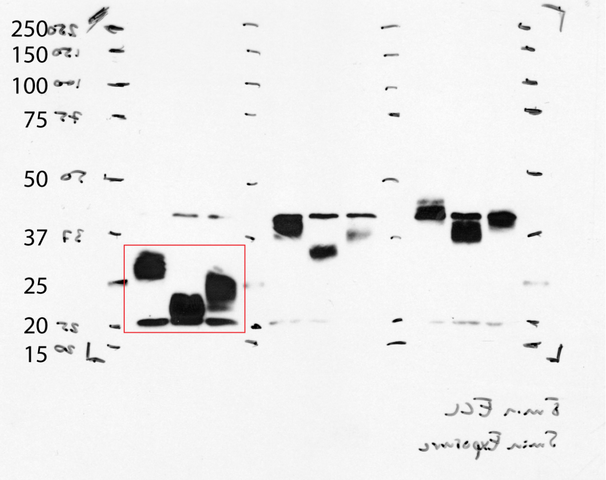


Figure 2B - RhoΔ38

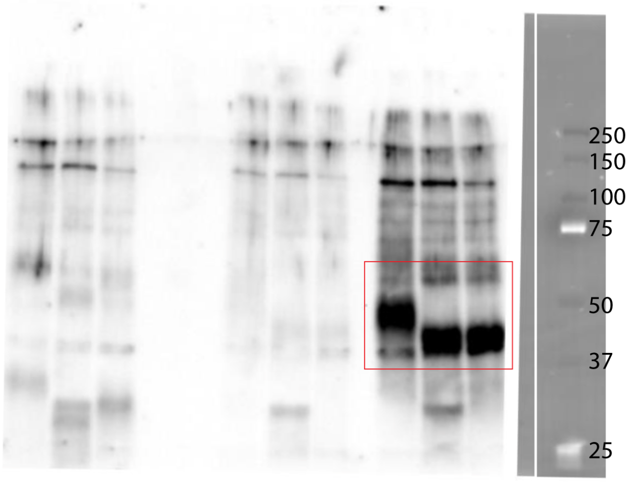

**Supplemental Table 1.** Summary of all transgenic constructs presented in this study

| Construct | Localization in WT | | OS extension in RhoKO | Figure |
|---|---|---|---|---|
| | OS | IS | | |
| FLAG-Rho-FL | + | - | + | Fig. 1 and Fig. 4 |
| EGFP-TM | + | + | NA | Fig. 1 and Fig. 4 |
| EGFP-Cb5-TM | - | + | NA | Fig. 1 |
| FLAG-RhoΔ5 | + | + | - | Fig. 2 and Fig. 4 |
| FLAG-RhoΔ25 | + | + | - | Fig. 2 and Fig. 4 |
| FLAG-RhoΔ38 | - | + | - | Fig. 2 and Fig. 4 |
| FLAG-TM-RhoCT | + | - | NA | Fig. 2 |
| FLAG-TM-RhoCTΔ5 | + | + | NA | Fig. 2 |
| FLAG-TM-RhoCTΔ25 | + | + | NA | Fig. 2 |
| FLAG-RhoΔ25+QVAPA | + | - | + | Fig. 3 and Fig. 4 |
| FLAG-RhoΔ38+QVAPA | - | + | - | Fig. 3 |
| FLAG-RhoΔ34 | - | + | NA | Fig. 3 |
| FLAG-RhoΔ38+KQVAPA | + | + | - | Fig. 3 and Fig. 4 |
| FLAG-RhoΔ25-CCAA+QVAPA | + | + | + | Fig. 4 |
| FLAG-RhoΔ25-CCAA | + | + | - | Sup Fig. 3 |
| FLAG-RhoΔ25-FRAA | - | + | - | Sup Fig. 3 |
| FLAG-RhoΔ25-FRAA+QVAPA | - | + | - | Sup Fig. 3 |
| FLAG-Rho-CC322,323AA | + | - | + | Fig. 5 |
| FLAG-Rho-FR313,314AA | + | + | + | Fig. 5 |
| FLAG-Rho-FRAA-CCAA | + | + | + | Fig. 5 |

“+” = attribute observed; “-” = attribute not observed
NA = not tested

**Supplemental Table 2.** Oligonucleotide primers to generate and sequence confirm constructs used in this study

| Primer Name | Sequence (5' --> 3') | Usage |
|---|---|---|
| pRho2.2K fwd seq | CGCCGCCGGGGATCCTCTAG | Amplify upstream from pRho-Rho; pRho sequencing |
| pRho2.2K rev seq | CCAGCCACCACCTTCTGATAG | Amplify downstream from pRho-Rho; pRho sequencing |
| Rho-Nterm-seq-R | TTCCGCCAGGTAGTACTGC | Sequencing mouse Rho |
| Rho-Mid-seqR | GATTCGTTGTTGACCTCAGG | Sequencing mouse Rho |
| SABRhoCtd5-NotI-BamHI-Rev | GCATGGATCCGCGGCCGCTTAGCTGGTCTCCGTCTTGG | Rhodopsin ∆5 truncation |
| Rho-D25-NotI-rev | ATGAGCGGCCGCTTAGCAGCACAGCGTGGTGAG | Rhodopsin ∆25 truncation |
| Rho1-314_NotI Rev | GCATGCGGCCGCTTACCGGAACTGCTTGTTCAAC | Rhodopsin ∆34 truncation |
| RhoD38_Rev | GCGGCCGCTCAGTTCAACATGATGTAGATGACC | Rhodopsin ∆38 truncation after $7^{th}$ TMD |
| RhoF313A_Fwd | CATGTTGAACAAGCAGGCCCGGAACTGTATGCTCACC | Mutagenesis primer for F313A mutant |
| RhoF313A_Rev | GGTGAGCATACAGTTCCGGGCCTGCTTGTTCAACATG | Mutagenesis primer for F313A mutant |
| RhoR314A_Fwd | CATGTTGAACAAGCAGTTCGCGAACTGTATGCTCACCACG | Mutagenesis primer for R314A mutant |
| RhoR314A_Rev | CGTGGTGAGCATACAGTTCGCGAACTGCTTGTTCAACATG | Mutagenesis primer for R314A mutant |
| NotI-+QVXPX-RhoD25_Rev | GCATGCGGCCGCTCAGGCTGGAGCCACCTGGCAGCACAGCG TGGTGAGCATAC | To append QVAPA motif to ∆25 constructs |
| Rho-CC322,323AA _Fwd | CTGTATGCTCACCACGCTGGCCGCCGGCAAGAATCCACT | Mutagenesis primer for CC322, 323AA |
| Rho-CC322,323AA _Rev | CAGTGGATTCTTGCCGGCGGCCAGCGTGGTGAGCATACA | Mutagenesis primer for CC322, 323AA |
| NotI-RhoD25-CCAA_Rev | GCATGCGGCCGCTCAGGCGGCCAGCGTGGTGAGCATACAG | Rhodopsin ∆25 truncation with CC322,323AA mutation |
| NotI-+QVXPX-RhoD25-CCAA_R | GCATGCGGCCGCTCAGGCTGGAGCCACCTGGGCGGCCAGC GTGGTGAGCATAC | ∆25 truncation with CCAA mutation and QVAPA appended |
| NotI-+QVAPA-RhoD38_Rev | GCATGCGGCCGCTCAGGCTGGAGCCACCTGGTTCAACATGA TGTAGATGACC | Rhodopsin ∆38 truncation with QVAPA appended |
| NotI-+KQVAPA-RhoD38_Rev | GCATGCGGCCGCTCAGGCTGGAGCCACCTGCTTGTTCAACAT GATGTAGATGA | Rhodopsin ∆38 truncation with KQVAPA appended |
| Flag-ActTM-Fwd | GACTACAAGGACGACGATGACAAGAGATCATTTCCGGAGATG | FLAG-tagged TMD fused to Rhodopsin C-termini |
| ActSS-Flag-Rev | CTTGTCATCGTCGTCCTTGTAGTCGCCAAGTATAGCACCTG | FLAG-tagged TMD fused to Rhodopsin C-termini |
